# Role of RNA Guanine Quadruplexes in Favoring the Dimerization of SARS Unique Domain in Coronaviruses

**DOI:** 10.1101/2020.04.07.029447

**Authors:** Cécilia Hognon, Tom Miclot, Cristina Garcia Iriepa, Antonio Francés-Monerris, Stephanie Grandemange, Alessio Terenzi, Marco Marazzi, Giampaolo Barone, Antonio Monari

**Affiliations:** Université de Lorraine and CNRS, LPCT UMR 7019, F-54000 Nancy, France; Department of Biological, Chemical and Pharmaceutical Sciences and Technologies, Università degli Studi di Palermo, Viale delle Scienze, 90128 Palermo, Italy; Department of Analytical Chemistry, Physical Chemistry and Chemical Engineering, Universidad de Alcalá, Ctra. Madrid-Barcelona, Km 33,600, 28871 Alcalá de Henares, Madrid, Spain; Chemical Research Institute “Andrés M. del Río” (IQAR), Universidad de Alcalá, 28871 Alcalá de Henares, Madrid, Spain; Departament de Química Física, Universitat de València, 46100 Burjassot, Spain; Université de Lorraine and CNRS, CRAN UMR 7039, F-54000 Nancy, France

**Keywords:** SARS, COVID-19, coronaviruses, protein-RNA interactions, molecular dynamics simulations, free energy profiles

## Abstract

Coronaviruses may produce severe acute respiratory syndrome (SARS). As a matter of fact, a new SARS-type virus, SARS-CoV-2, is responsible of a global pandemic in 2020 with unprecedented sanitary and economic consequences for most countries. In the present contribution we study, by all-atom equilibrium and enhanced sampling molecular dynamics simulations, the interaction between the SARS Unique Domain and RNA guanine quadruplexes, a process involved in eluding the defensive response of the host thus favoring viral infection of human cells. Our results evidence two stable binding modes involving an interaction site spanning either the protein dimer interface or only one monomer. The free energy profile unequivocally points to the dimer mode as the thermodynamically favored one. The effect of these binding modes in stabilizing the protein dimer was also assessed, being related to its biological role in assisting SARS viruses to bypass the host protective response. This work also constitutes a first step of the possible rational design of efficient therapeutic agents aiming at perturbing the interaction between SARS Unique Domain and guanine quadruplexes, hence enhancing the host defenses against the virus.

**TOC GRAPHICS:** 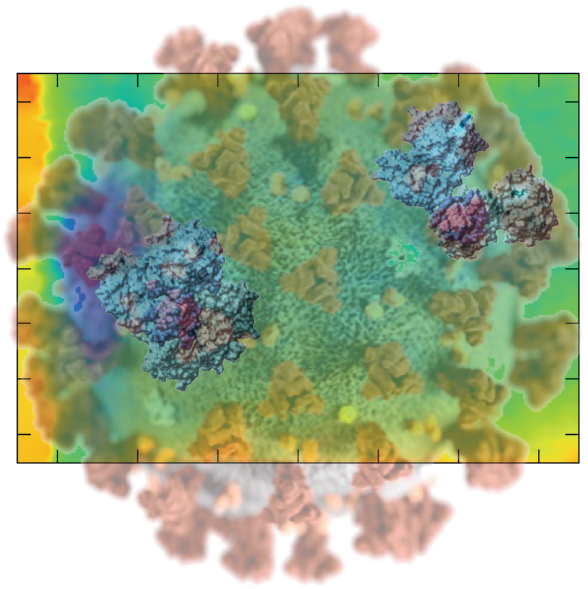

---

Coronaviruses inducing severe acute respiratory syndrome (SARS) gained first worldwide attention in 2003 following the outbreak of SARS disease that mainly affected different south-eastern Asian countries.^1–3^ By the end of 2019 and the beginning of 2020, a novel coronavirus, styled SARS-CoV-2, has emerged and caused the Covid-19 pandemic.^4–7^ At the time of writing this communication, Covid-19 has spread in all continents, excluding Antarctica. In April, the 5^th^ 2020 a number of more than 1 million confirmed cases has been reported, as well as more than 66,000 deaths.^8^ Furthermore, severe limitations and social distancing measures have been implemented by an increasing number of countries, in the attempt to limit the spreading of the pandemic.^9^ Indeed, coronaviruses, and SARS-CoV-2 in particular, are characterized by high infectious and transmissibility rates, and may, in certain cases, severely attack the lungs and the respiratory system, an occurrence which can lead to the necessity of respiratory assistance and even to death, especially in presence of others comorbidities.^10–12^

Since the first outbreak of the pandemic, important scientific works appeared, aiming at understanding at a molecular level the mechanisms of action of SARS-CoV-2, and corona viruses in general, that may be related to its transmissibility and mortality. Important successes have notably been achieved in both genome sequencing and structural resolution of the different viral proteins, also with the assistance of molecular modeling and simulation.^13–21^ In fact, the possibility of finding efficient therapeutic strategies requires the elucidation of the fine mechanism that mediate the virus attack to the host cell, the resistance to the host immune system, and finally its virulence.^22–25^

Among the multifaced protein envelope of coronaviruses the so-called non-structural protein 3 (Nsp3) represents a large protein, localized in the cytoplasm, consisting of about 2,000 amino acids and comprising several different domains, also including a transmembrane region. Among the different domains of Nsp3,^26,27^ whose precise function of some of them has not been entirely clarified yet, the so-called SARS Unique Domain (SUD) deserves a special attention, since it is present only in SARS-type coronaviruses and hence it has been associated to the increased pathogenicity of this viral family.

The structure of SUD (presumably a common domain of different SARS viruses) has been resolved experimentally,^28–30^ and it has been proved that the macrodomain is indeed constituted by a dimer of two symmetric monomers. Furthermore, both experimental and molecular docking investigations have pointed out a possible favorable interaction of SUD with nucleic acids, and in particular with RNA in G-quadruplex (G4) conformation.^28^ The presence of a high density of lysine residues at the interface between two SUD monomers, forming a positively charged pocket, also suggests that RNA may be instrumental in favoring SUD dimerization, due to the negative charge of the RNA backbone hence suggesting the occurrence of electrostatic attraction. This observation may have a highly important biological implication since the dimerization has also been connected to the SUD native function. Tan *et* al^28^ have proposed that the ability of SUD to recognize and bind specific viral and/or host G4 sequences may have implications in regulating viral replication and/or hampering the host response to viral infection, as schematized in Figure 1. The hypothesis is based on the identification of G4 sequences in key host mRNA that encode proteins involved in different signaling pathways such as apoptoting or survival signaling.^31–38^ These proteins could induce a controlled cellular death of infected cells slowing down or stopping the infection, or promote cell survival to delay apoptosis by producing antiviral cytokines.^37^ However, the removal of the mRNA necessary to produce these signaling factors by viral SUD may impair the apoptosis/survival response pathways allowing massive cell infection.^28,37^

**Figure 1.**
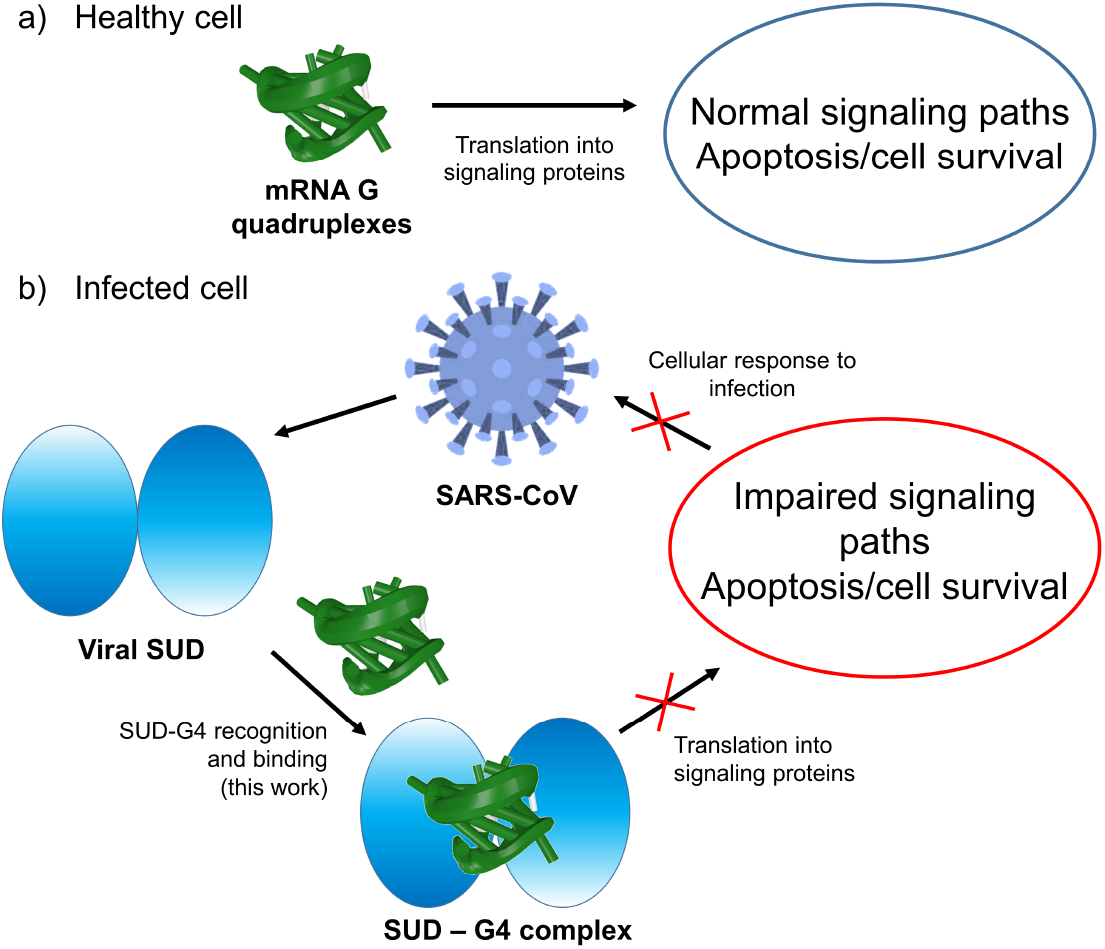
Schematic representation of the mRNA function in a) a healthy cell and b) an infected cell by coronavirus. Panel b) showcases the influence of viral SUD binding to G4 sequences of mRNA that encodes crucial proteins for the apoptosis/cell survival regulation and other signaling paths.

In this letter, we report an extended all-atom molecular dynamics (MD) study of the interactions produced between a dimeric SUD domain and a short RNA G4 sequence. The crystal structure of the protein (pdb 2W2G) and of the oligonucleotide (pdb 1J8G)^39^ have been chosen coherently with the experimental work performed by Tan et al.^28^ Even though the chosen SUD starting structure belongs to the 2009 SARS-CoV, the very high nucleotide affinity^40^ and the global conservation of the Nsp protein suggest that the RNA binding spots should be globally preserved. This is also further justified by the fact that SARS-CoV-2 Nsp has also been recognized to suppress host gene expression and hence inhibit the immune response.^41^ Equilibrium MD has allowed to assess some of the hypothesized complexation modes between G4 and SUD, while also highlighting the most important interactions patterns at an atomistic level, and the effects of G4 in maintaining the dimer stability. Furthermore, to better sample the multidimensional conformational space and to quantify the strength of the interactions coming into play, the free-energy surface has been explored using enhanced sampling methods. A two-dimensional (2D) free energy profile has been computed along two coordinates defining the distance between the centers of mass of G4 and one SUD domain (G4-SUD_A_), and the two SUD domains (SUD_A_-SUD_B_), respectively. The corresponding 2D potential of mean force (PMF) was obtained using a recently developed combination of extended adaptative biased force (eABF)^42^ and metadynamics,^43^ hereafter named meta-eABF.^44,45^ Both protein and RNA have been described with the amber force field^46^ including the bsc1 corrections,^47,48^ and the MD simulations have been performed in the constant pressure and temperature ensemble (NPT) at 300K and 1 atm. All MD simulations have been performed using the NAMD code^49^ and analyzed via VMD,^50^ the G4 structure has also been analyzed with the 3DNA suite.^51,52^ More details on the simulation protocol can be found in Supplementary Information (SI). To obtain starting conformations, the RNA was manually positioned in two different orientations close to the experimentally suggested SUD interaction area.^28^ The equilibrium MD evolved yielding two distinct interaction modes, as reported in Figure 2. In particular, we can easily distinguish between a first mode of binding in which the G4 mainly interacts with only one SUD monomer, called monomeric binding mode, and a second one in which the nucleic acid is firmly placed at the interface between the two protein monomeric subunits, referred as dimeric binding mode. Note that while for the monomeric mode we easily found a suitable starting point, two independent 200 ns MD trajectory were run to characterize the dimeric mode.

**Figure 2.**
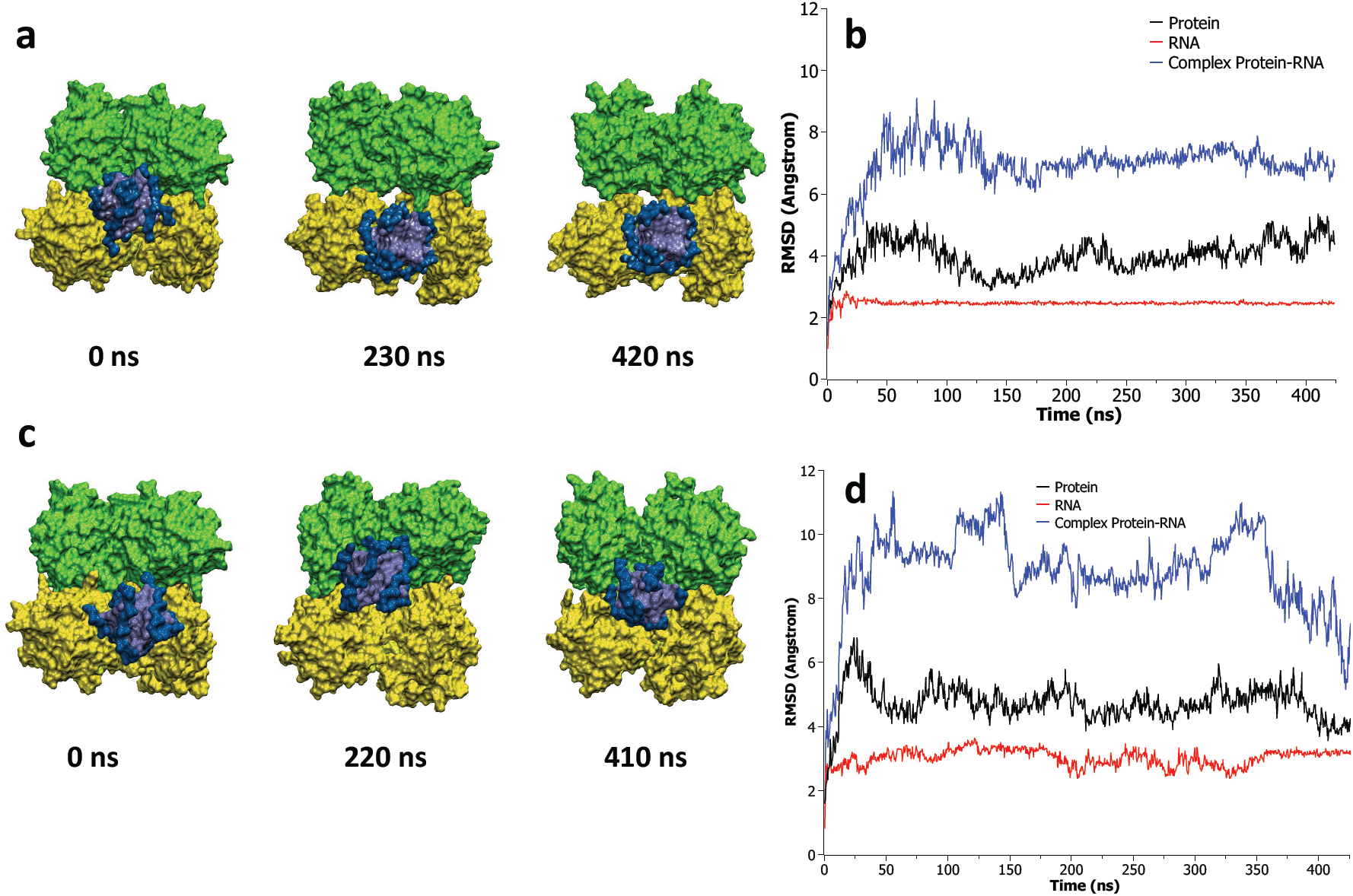
Representative snapshots of the SUD/G4 complex as extracted from the equilibrium MD simulation for the monomeric interaction mode (a) together with the corresponding RMSD for the protein and RNA fragments (b). Representative snapshots (c) and corresponding RMSD time evolutions (d) for the dimeric interaction mode. Note that the partial RMSD time series have been calculated aligning the system to the RNA or the protein, respectively.

The corresponding root mean square deviations (RMSD) with respect to the initial structure are also reported and globally show that both the RNA and the protein units are stable. As expected, a slightly larger value of the RMSD is observed for the protein, as a consequence of its larger flexibility compared to the rigid G4 structures (Figure 2 d, c). Note also that the slight initial increase of the protein RMSD observed for the dimeric mode is due to the necessity of a slight structural rearrangement to accommodate the G4 in the interaction pocket. Both modes are globally stable all along the MD trajectory, and no spontaneous release of the G4 is observed. At the end of the MD trajectory the dimeric mode exhibits a slight slipping toward the center of the interface area. This is probably due to the relative short length of the oligomer, also in agreement with previous experimental observations.^28^

Interestingly, the specific interaction patterns between G4 and the protein are different between the two binding modes, providing an extremely important different behavior. Indeed, the main driving force for the dimeric-like binding mode is the presence of extended electrostatic interactions between the negatively charged RNA backbone and the highly positive interaction pockets of the SUD complex. As a matter of fact, eleven Lys residues in the interaction pocket (shown in green in Figure 3a) allowing persistent salt-bridges and hydrogen-bonds with the RNA phosphates. This finding is evidenced by the radial distribution function (RDF) between these positively charged lysine side chains and the negatively charged phosphate oxygen atoms of G4 (depicted in dark blue in Figure 3b), which shows a very intense and sharp peak at around 2 Å (Figure 3a). Interestingly, a secondary peak in the RDF is also observed at 3.5 Å, probably defining a second layer of interaction patterns that should contribute to the overall stabilization of the binding. Conversely, the monomeric mode is driven by interactions mainly involving the terminal uracil moieties and the top guanine leaflet instead of the phosphate backbone of G4. As shown in Figure 3b, hydrophobic interactions emerge through the action of a triad of amino acids, namely Ser236, Leu237, and Asn238, that position themselves on top of the terminal leaflet also interacting with the peripheral uracil nucleobases, in a mode strongly resembling the top-binding experienced by a number of G4 drugs.^53–55^ This is nicely confirmed by the analysis of the time series of the distance between the α–carbon of these amino acids and the nearby guanine that readily drops at around 5 Å and stays remarkably stable all along the MD. Interestingly, the interaction is sufficiently strong to induce a partial deformation of the planarity of the G4 leaflets. Even though from considerations based on chemical intuition those interactions could be referred as mainly due to dispersion, the inherent parameterization of the amber force field does not allow to completely disentangle and decompose the polarization, dispersion and electrostatic contributions. The fact that the monomeric binding mode is driven by non-covalent interactions with one of the exposed G4 leaflets may also contribute explaining the fact that longer G4 sequences are preferentially recognized by the SUD interface region. Indeed, in this case, for obvious statistic reasons, the ratio between the interaction with the backbone or with the terminal leaflet clearly favors the former. On the other hand, this mode may also act efficiently in the process of recruitment of RNA, either viral or cellular, efficiently anchoring the oligomer that can subsequently be safely displaced through the interface area.

**Figure 3.**
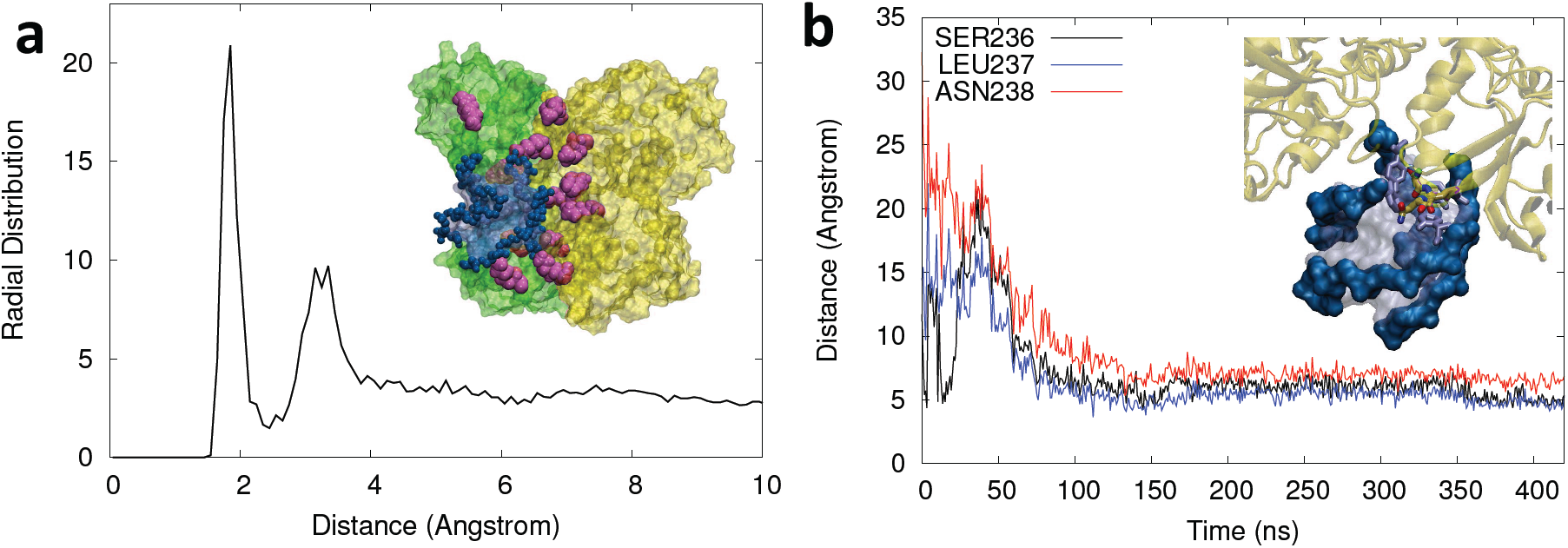
(a) RDF between the RNA phosphate oxygens and the Lys NH3 hydrogens for the dimeric interaction mode (see figure 2c). The inlay shows a representative snapshot showing the network of Lys (in purple, van der Waals representation) in the interaction pocket surrounding the RNA G4, whose backbone phosphate groups are highlighted in dark blue. (b) Time series of the distances between the α-carbon of Ser236, Leu237, and Asn238 and the G4 nearby uracil or guanine oxygen atom. A zoom of the representative snapshot (Figure 2a) showing the corresponding interactions is also provided.

Apart from the different positioning of the G4, other structural evidences can already be surmised from the visual inspection of the MD trajectory. In particular, the SUD dimers appear more compact and the interface region better conserved when the RNA G4 adopts the dimeric binding mode, as can also been appreciated in Figure 4. These results clearly indicate that the dimeric mode leads to a greater stability of the G4-SUD complex.

**Figure 4.**
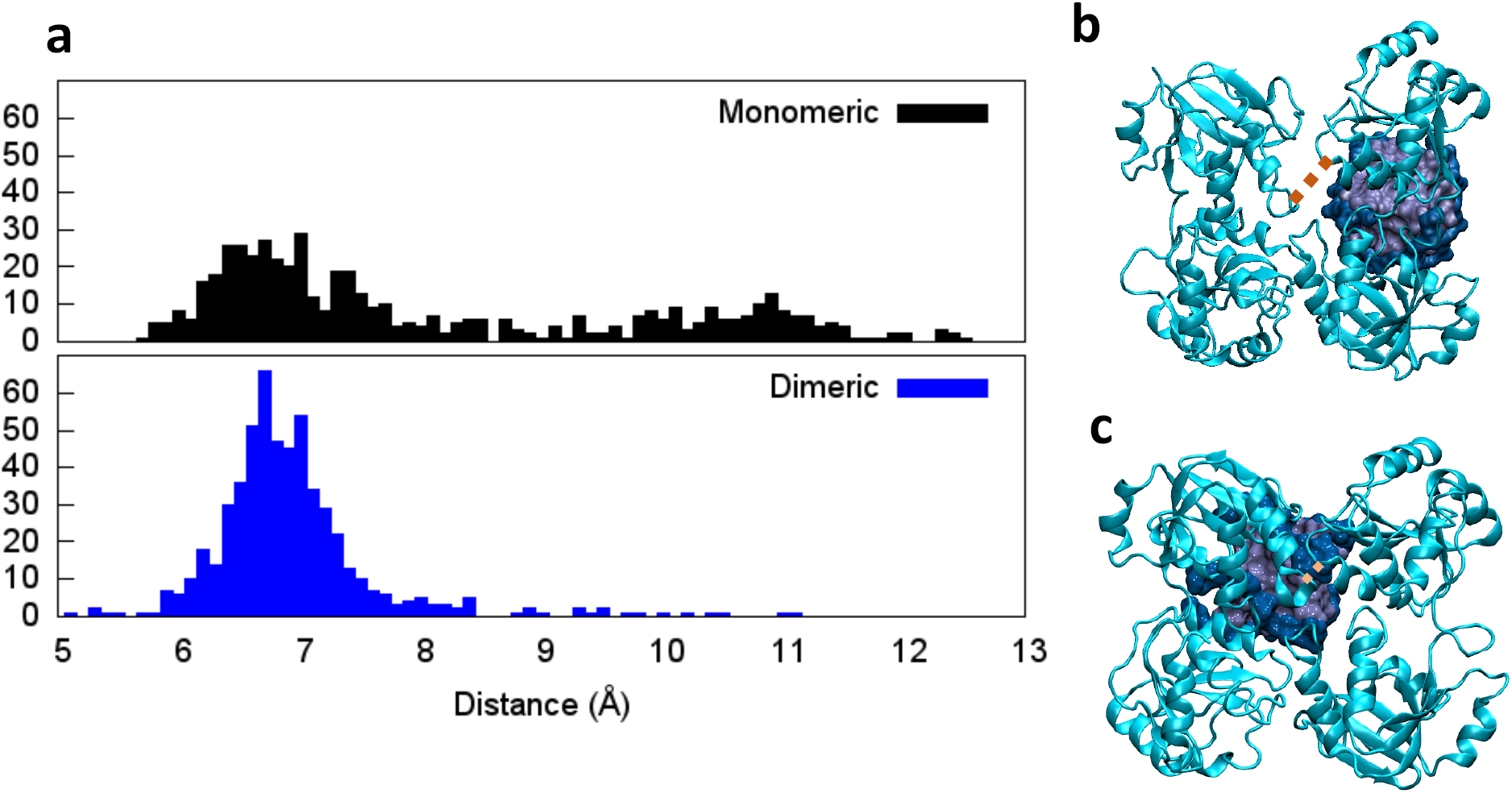
Distribution of the distances between the α-carbon of Arg266 and Ala366 (a) forming a tweezer in the SUD subdomains interface region, representative snapshot showing this region for the monomer (b) and the dimer (c) conformation. The dashed orange line indicates the position of the protein residues corresponding to the distance measured.

Indeed, the presence of G4 in the dimeric binding mode induces rigidity in the SUD subunits interface due to the formation of the dimer. This effect can be quantified by analyzing the distribution of the distance of a tweezer in the SUD interface area formed by the residues Ala355 and Arg266, as reported in Figure 4. Indeed, while in the case of the dimeric-like conformation a peaked distribution centered at relative close distances (6.7 Å) is observed, indicating a closed and quite rigid disposition, a much broader and bimodal distribution is found for the monomer-like conformation, presenting, most notably, a secondary maximum at about 11 Å, which confirms the partial destabilization of the SUD subdomains interface and the greater flexibility induced by this binding mode.

To further examine the conformational space spanned by the G4/SUD complex, and in particular the role of the RNA in favoring the dimerization and the structure of the interface, we resorted to enhanced sampling MD simulations to obtain the 2D free energy profile along two relevant collective variables: first, the distance between G4 and SUD, and second, the separation between the two SUD subdomains. Our choice of collective variables does not allow to explore the binding between the two surfaces of the SUD domain, however from the results of Tan et al.^28^ it is clear that the interaction with RNA takes place preferentially at the positively charged interface. On the other hand, our methodology is perfectly adapted to characterize the influence of the binding of RNA to the stabilization of the interface between the two SUD monomers since it allows the sliding of the G4 on the SUD surface. The PMF is reported in Figure 5 together with representative snapshots along the reaction coordinates. From the analysis of the PMF, one can evidence the presence of an evident minimum in the free energy profile corresponding to the situation in which the G4 is interacting through the dimer mode, in which the SUD dimer is compact (Figure 5b). The free-energy stabilization, with respect to the situation in which the G4 is well separated from the protein, amounts to about 6 kcal/mol. Around the principal minimum a slightly less stable and extended region is also observed having a stabilization free energy of about 3-4 kcal/mol and corresponding to the sliding of G4 in the monomer conformation (Figure 5c). The rest of the free energy surface appears instead rather flat, and no appreciable barriers are observed along the collective variable. The topology of the free energy surface hence accounts for the possibility to observe important conformational movements, leading to open conformations in which the SUD subdomain interface has been basically destroyed (Figure 5d). However, such conformations are instead hampered by the dimer-like conformation of the RNA. The free energy map unambiguously shows that the dimer mode is the preferred one, and also confirms the role of the G4 binding in maintaining the dimeric SUD conformation, since no appreciable free energy barrier for the opening of the SUD dimer is observed when the RNA is unbound. Thus, the dimer mode binding site clearly constitutes a specific target that may help in the development of new efficient antiviral agents against coronaviruses.

**Figure 5.**
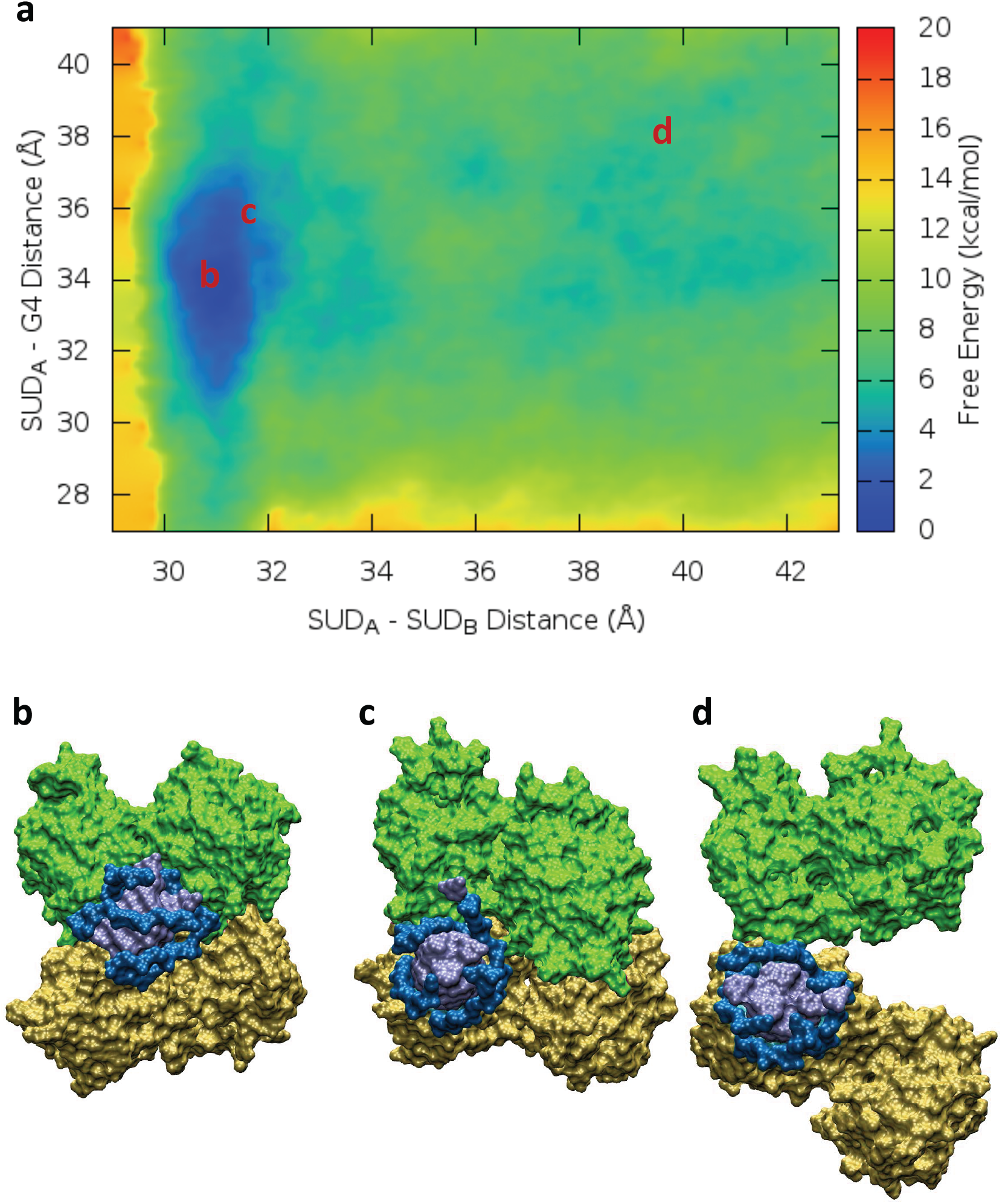
a) 2D Free energy profile describing the interaction with RNA and SUD dimerization. Representative snapshots are also provided describing the principal minimum (b), the secondary minimum (c), and an open SUD conformation (d), the position of the selected snapshots on the PMF map is also indicated in red.

By using a combination of equilibrium and enhanced sampling MD, we have identified the structure and dynamics of the viral SUD dimers and their interaction with RNA oligomers forming guanine quadruplexes. Even if the accuracy of the obtained results is obviously dependent on the choice of the specific force field and of the specific collective variables, we believe we have been able to offer a significant sampling of the binding between SUD and RNA G4, and of its effects on the stabilization of the SUD complex and the monomer-monomer interface. In particular, we have clearly evidenced the existence of two different interaction modes, one happening at the subdomains interface (dimeric binding mode) and another involving only one of the SUD monomers. While the former mode is mostly driven by electrostatic interactions and appears to be more stable and is essential in rigidifying the protein dimer and stabilizing the SUD-G4 interface, the latter assures a great flexibility to the protein and is guided by non-covalent interactions. Such stable interaction between SUD and G4-RNA can be related to the translational default of the concerned proteins (see Figure 1). Furthermore, the monomer interaction mode, happening in a more solvent exposed region of the protein, can also play a role in recruiting RNA fragments in the first step of the SUD-G4 recognition.

SUD dimer is thought to have an important biological role in allowing coronaviruses to bypass host cell protective and immune systems, hence allowing the virus morbidity and transmissibility. Indeed, and even if in the present work we used a protein belonging to SARS-CoV, we believe that the mechanism underlined is quite general and may represent one of the reasons of the high pathogenicity of SARS agents. As a matter of fact, the role of SARS-CoV-2 Nsp in suppressing host gene expression as also been recently evidenced.^41^ The destabilization of the SUD/RNA complex, that could be achieved for instance by specific G4 ligands, or by competitors for the two G4 recognition spots evidenced in SUD, may constitute appealing potential therapeutic strategies.

This is particularly true since these protein domains are shared by the family of SARS-type viruses and thus represent a molecular target that could stay operative in case of future mutations leading to new viral strains. As a continuation of the present work, we plan to address the study of the interaction between specific ligands and RNA G4 oligomers, also taking into account the perturbation induced on the recognition by SUD.

## ASSOCIATED CONTENT

### Supporting Information

Details of the protocol used for equilibrium MD and free energy determination, analysis of the structural parameters of the G4 unit and of the set-up of the SUD initial structure, time series of the distance between the residues defining the opening of SUD. The following files are available free of charge.

SI_SUD.pdf (PDF)

## AUTHOR INFORMATION

The authors declare no competing financial interests.

## ACKNOWLEDGMENT

Support from the Université de Lorraine and CNRS is gratefully acknowledged. T.M. thanks University of Palermo for funding a joint Ph.D. program A.F.-M. is grateful to Generalitat Valenciana and the European Social Fund for a postdoctoral contract (APOSTD/2019/149) and the *Ministerio de Ciencia e Innovación* (project (CTQ2017-87054-C2-2-P) for financial support. French CNRS and IDRIS national computing center are acknowledged for graciously providing access to computational resources in the framework of the special Covid-19 mobilization under the project “Seek&Destroy”. Part of the calculations have been performed on the LPCT local computing cluster and on the regional ExpLor center in the frame of the project “Dancing Under the Light”.

