## Supplementary Information for "Role of RNA Guanine Quadruplexes in Favoring the Dimerization of SARS Unique Domain in Coronaviruses"

**Details of the Equilibrium MD Simulation**

All structures involving the formation of a complex between SUD and RNA were created in silico with the PyMol software. SUD initial structure has been obtained from the 2W2G pdb entry while the G-quadruplex RNA from the 1J8G entry. The RNA was manually positioned in two different orientations close to the experimentally suggested interaction area. The initial structures were solvated in a TIP3[1] cubic water box having a size of 104 Å with ~ 31 000 water molecules. 15 $K^+$ cations were added to assure neutrality. All the molecular dynamic simulations have been performed using NAMD 2.13 code[2] with amber force field parm99[3] force field including BSC1[4] correction for nucleic acids. To increase the performance, we used H Mass Repartition technique[5] which consist to fix the mass of H from 1.008 to 3.024 Da. In this way, a time step of 4 fs was used to integrate the Newton equation of motion in combination with the Rattle and Shake algorithms[6]. The cut off for non-covalent interactions was set to 9 Å while the particle mesh Ewald procedure (PME)[7] was consistently used to calculate electrostatic interactions.

The simulation protocol used has been the following: 1000 steps of minimization have been applied to remove bad contacts, followed by a first dynamic of 36 ns in which constraints imposed on the protein and RNA atoms were progressively released. The MD simulations of 425 ns have been performed in the NPT ensemble at a temperature of 300k and the pressure of 1 atm. Isothermal and isobar conditions were maintained using the Langevin thermostat and piston, periodic boundary conditions (PBC) were consistently used.

The trajectories have been analyzed and representative snapshots rendered using VMD program[8].

**Topology of the RNA G-quadruplex**

The selected RNA is composed of four RNA strands of sequence 5'-UGGGGU-3' that fold in a G4 arrangement (Figure S1) stabilized by bivalent $Sr^{2+}$ cations, the analysis of the topology of the RNA, also performed using x3DNA code confirmed that the strand forms a stable G4 assuming a parallel arrangement and a right-hand twist of 28.7°, all the guanine leaflet maintained a remarkable planarity. In addition to the formation of the guanine tetrad, a rther unusual uracil tetrad can be observed at the 3'-end side of the G4.

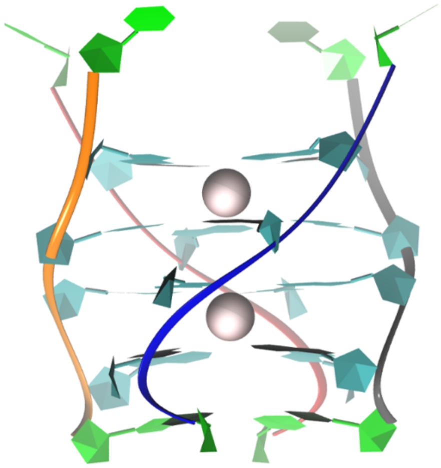

Figure S1) Representation of the RNA G4 used in the present work, the terminal uracil nucleobases are highlighted in green.

**Topology of the SUD core**

SUD domain is derived from the part of the Nsp3 protein. It is a homodimer whose sequence is fully known; however, some residues were missing from the crystal structure, and have been added with the pymol utilities (Figure S2) showing a good agreement. Hydrogen atoms have been added, considering the pKa of the involved aminoacids and a disulfur bridge between two cysteine residues tethering the two monomers has been added.

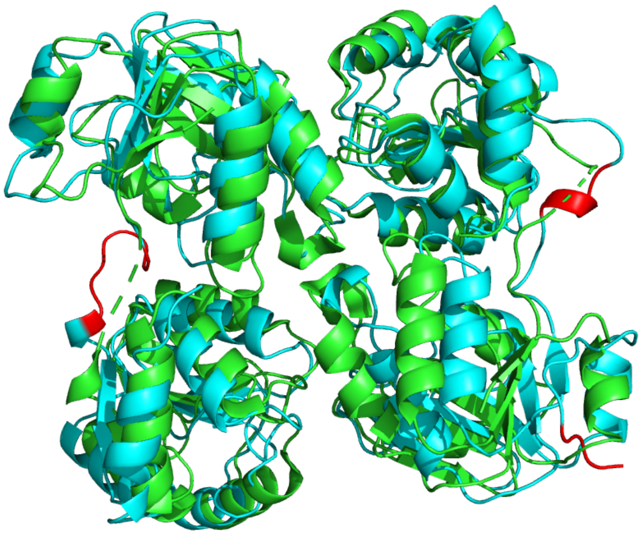

Figure S2) Alignment between the crystal structure of the SUD domain and the initial topology the added residues are represented in red

**Details of the Free Energy Determination**

The 2D free energy profile for the characterization of the SUD/G4 complex was performed sampling the collective variable defined by the variation of the distance between the two monomers (a in figure S3); and the distance between monomer 1 and DNA (b in Figure S3) with respect to the initial structure. The potential of mean force (PMF) profiles have been obtained using the combination of the extended adaptive biasing force (eABF) and metadynamics algorithms (meta-eABF). A harmonic bias potential was used. The free energy profile was estimated in the distance interval comprised between 29 Å and 43 Å for a; 27 Å and 41 Å for b in a single window. The instantaneous force values were accumulated in bins of 0.1 Å. The lower and upper walls (boundary potentials) were constrained with a constant force of 100 kcal/mol.

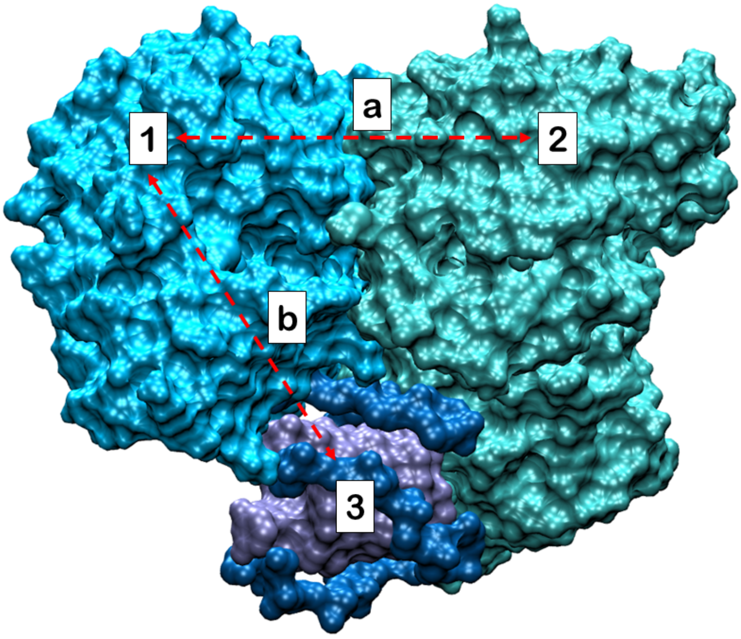


Figure S3: Definition of collective variable used for the 2D free energy profile.

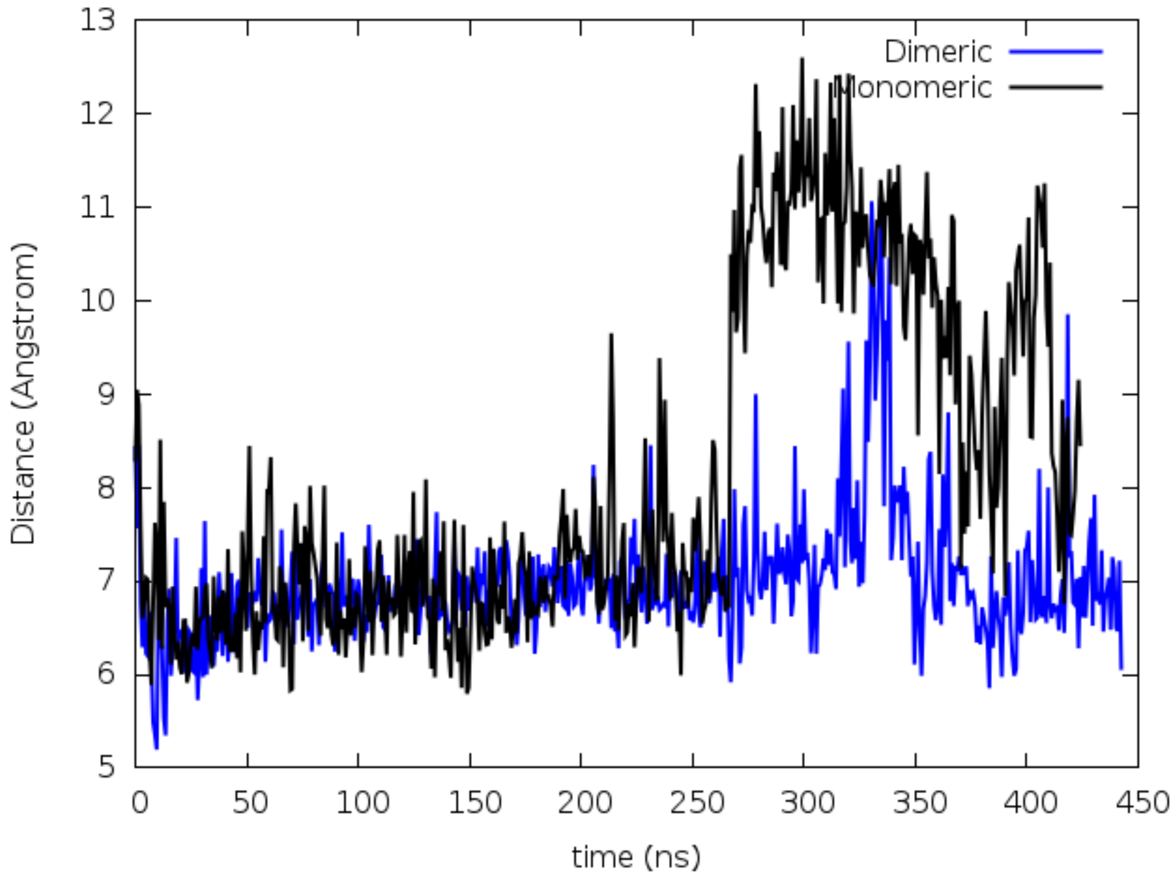


Figure S4) Time series of the distances between the □-carbon of Arg266 and Ala366 forming a tweezer in the SUD subdomains interface region.
